# Inefficient ZAP70-Signaling Blunts Antigen Detection by CAR-T-Cells

**DOI:** 10.1101/720417

**Authors:** Venugopal Gudipati, Julian Rydzek, Iago Doel Perez, Lydia Scharf, Sebastian Königsberger, Hermann Einsele, Hannes Stockinger, Michael Hudecek, Johannes B. Huppa

**Author notes:** Authors contributed equally to this study (Venugopal Gudipati, Julian Rydzek). Division of Clinical Microbiology, Department of Laboratory Medicine, Karolinska Institutet, Karolinska University Hospital Huddinge, Stockholm, Sweden (Lydia Scharf). Eurofins BioPharma Product Testing Munich GmbH, Behringstraße 6, D-82152 Planegg, Germany (Sebastian Königsberger).

## Abstract

Rational design of chimeric antigen receptors (CARs) with optimized anti-cancer performance mandates detailed knowledge of how CARs engage tumor antigens and how antigen-engagement triggers activation. We analyzed CAR-mediated antigen recognition via quantitative single molecule live-cell imaging and found the sensitivity of CAR-T-cells towards antigen approximately 1000-times reduced when compared to T-cell antigen receptor (TCR)-mediated recognition of nominal peptide/MHC complexes. While CARs outperformed TCRs with regard to antigen binding within the immunological synapse, proximal signaling was significantly attenuated due to inefficient recruitment of the tyrosine-kinase ZAP70 to ligated CARs and its reduced concomitant activation and subsequent release. Our study exposes signaling deficiencies of state-of-the-art CAR-designs, which limit at present the efficacy of CAR-T-cell therapies to target tumors with diminished antigen expression.

## INTRODUCTION

Adoptive immunotherapy employing chimeric antigen receptor (CAR)-modified T-cells has shown considerable potential as an effective treatment option for advanced B-cell malignancies (Maude et al., 2018; Maus et al., 2014; Park et al., 2018). CARs have been empirically designed to emulate the antigen-binding properties of monoclonal antibodies (mAb) and the signaling properties of the T-cell antigen receptor (TCR) as well as co-stimulatory receptors. CARs typically feature an extracellular, mAb-derived single-chain variable fragment (scF_V_) targeting the tumor associated antigen (TAA), which is linked to a spacer and transmembrane domain as well as intracellular signaling modules comprising domains from CD3ζ and co-stimulatory molecules such as CD28, 4-1BB and others (Finney et al., 1998; Imai et al., 2004; Maher et al., 2002). While this modular, rather one-dimensional architecture affords much flexibility in clinical applications, it conceivably limits the extent to which CARs reproduce the complexities of the signaling responses, which have coevolved with TCRs and costimulatory receptors (Srivastava and Riddell, 2015).

Of note, T-cells respond to the presence of even a single nominal peptide/MHC complex (pMHC) on the surface of antigen presenting cells (Huang et al., 2013; Irvine et al., 2002; Purbhoo et al., 2004). This remarkable capacity of T-cells is in stark contrast to the observation that more than 20% of patients suffer from cancer relapses after CD19-CAR-T-cell therapy due to the emergence of tumors which express antigen at low or undetectable levels (Maude et al., 2014; Maude et al., 2018; Turtle et al., 2016). The extent to which CD19 loss is partial or complete remains to be resolved, but reduced surface expression as widely regarded a major concern after CD19-CAR therapy (Orlando et al., 2018; Sotillo et al., 2015). In fact, more than 50% of the patients who had attained complete remission in a recent CD22-CAR clinical trial, relapsed later due to diminished cell surface CD22 expression (Fry et al., 2018). Addressing these complications in a comprehensive manner mandates a deep understanding of how CARs act on the molecular level, and more specifically, how synaptic antigen engagement triggers ensuing intracellular signaling (Srivastava and Riddell, 2015).

A major challenge in studying CAR-mediated antigen recognition and early signaling is rooted in the fact that underlying mechanisms involve short-lived protein-protein and protein-lipid interactions within highly specific cellular compartments (Baumgart and Schütz, 2015; Lillemeier et al., 2010). Conventional biochemical methods require, without exception, cell disruption and hence do not account for geometrical constraints and nonlinear properties of the immune synapse (Huppa and Davis, 2013). We have therefore devised a minimally invasive live-cell imaging methodology, which allows for quantitation of synaptic antigen-binding and ensuing downstream signaling with single molecule resolution (Axmann et al., 2015).

Following this approach, we found that virus-specific cytolytic T-cells (CTLs) expressing a CAR required approximately 1000-times more antigen to initiate intracellular signaling their CAR than via their endogenous TCR. Limitations in antigen detection were largely independent of the nature of the targeted antigen, the position of the epitope within the antigen and the presence of CD28- or 4-1BB-derived costimulatory signaling modules within the CAR. Furthermore, CAR surface expression levels were not functionally limiting, as synaptic CAR-antigen engagement outperformed synaptic TCR-antigen binding at lower antigen densities, at which TCR-signaling was still robust but CAR-signaling was no longer maintained. Instead, we witnessed poor recruitment of the zeta-chain-associated protein kinase 70 (ZAP70) to ligated CARs as well as attenuated concomitant ZAP70-activation. We hence conclude that inefficient triggering mechanisms, which link extracellular CAR engagement to ZAP70-activation, limit the capacity of current CAR-T-cells to target tumor cells expressing TAAs at low levels.

## RESULTS

### Imaging platform for comprehensive analysis of antigen recognition by CAR-T-cells

Efforts to quantitate receptor-antigen interactions and ensuing signaling events at the molecular level within conjugates formed between T-cells and antigen presenting cells (APCs) are typically challenged by three-dimensional cellular movement, phototoxicity owing to high-powered sample excitation and elevated noise levels arising from light scattering and cellular background fluorescence. To sidestep these issues altogether, we customized a planar glass-supported lipid bilayer (SLB)-based system to serve as surrogate APCs and to allow for imaging in Total Internal Reflection (TIR) mode (Figure 1A). Evanescent light waves, which result from the totally reflected laser beam, penetrate only 100 to 200 nanometers into the SLB-engaged T-cell (Axelrod et al., 1984). This reduces phototoxicity and cellular background to levels that are adequate for visualizing and quantitating plasma membrane proximal events with single molecule resolution without affecting cellular integrity (Douglass and Vale, 2005; Huppa et al., 2010).

**Figure 1:**
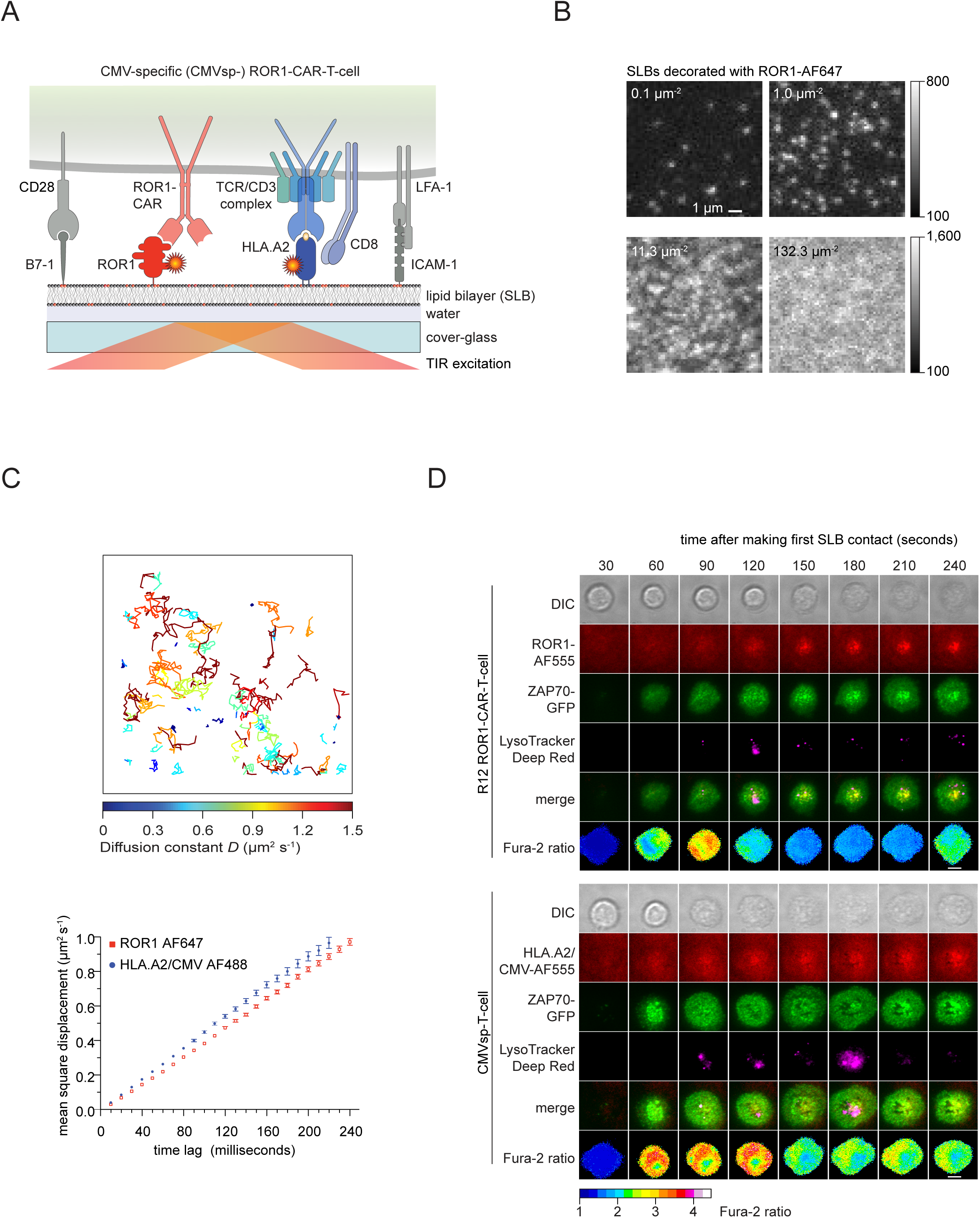
Protein-functionalized glass-<u>s</u>upported lipid <u>b</u>ilayer (SLB) system allows for highly resolved spatiotemporal imaging of CAR or TCR mediated antigen recognition. **(A)** Schematic representation of an SLB featuring fluorescently-labeled antigens (HLA.A2/CMV or ROR1), the adhesion molecule ICAM-1 and the costimulatory molecule B7-1 for recognition by CAR-T-cells. Extracellular portions of the proteins of interest are extended with a poly-histidine (12xHis) tag to interact with 18:1 DGS-NTA(Ni) (indicated with red headgroups) present in the SLB. **(B)** Micrographs of SLBs featuring ROR1-AF555 at indicated densities (0.1 to 132.3 ROR1 molecules µm^-2^). Note diffraction-limited fluorescence events appearing in the upper two micrographs reflecting single molecules. (**C**, upper panel) Representative traces of single ROR1-AF555 molecules diffusing across the SLB. **(C**, bottom panel) The rate of the lateral diffusion of SLB-anchored proteins was derived by plotting the mean square displacement (MSD) as a function of the time-lag. The linear increase in MSD over time indicates unconfined diffusion with the diffusion coefficient *D* being proportional to the plotted slope (*MSD* = 4 *D t*, with *D* = diffusion constant and *t* = time; n ≥ 500 per molecular species, error bars represent standard deviation). (**D**) Time-lapse microscopy to monitor antigen recognition by ROR1-transduced CMVsp-T-cells confronted with SLBs featuring ICAM-1, B7-1 as well as HLA.A2/CMV or ROR1. Enrichment of antigen underneath the T-cells is reflective of antigen engagement and occurred as soon as T-cells contacted SLBs. (CAR-) T-cell activation became evident through (i) the synaptic recruitment of lentivirally expressed ZAP70-GFP to engaged antigen and (ii) a sizable increase in intracellular calcium as monitored with the ratiometric calcium-sensitive dye Fura-2. Lytic granules stained with LysoTracker Deep Red moved concomitantly to the center of immunological synapse and into the TIR illumination field to initiate target cell killing. Shown images are representative of more than 30 T-cells picked at random and derived from two different donors (scale bar; 5µm)

Throughout this study we incorporated the synthetic lipid 18:1 DGS-NTA(Ni) into SLBs to provide acceptor sites for poly-histidine-tagged recombinant proteins (for details refer to Methods section). These included the receptor tyrosine kinase-like orphan receptor 1 (ROR1) for recognition by ROR1-specific CAR-T-cells, the adhesion molecule ICAM-1 binding to the integrin LFA-1 and the co-stimulatory molecule B7-1 (Figure 1A, B, Figure S1A). For stimulation of cytomegalovirus (CMV) -specific (CMVsp) CTLs, which served as a reference for CAR-T-cell performance, ROR1 was substituted with the MHC class I molecule HLA-A*0201 loaded with antigenic CMV-derived peptide pp65 (HLA.A2/CMV). Lateral mobility of SLB-associated-proteins was verified via fluorescence recovery after photobleaching (FRAP) and single dye tracing (SDT): more than 90% of ROR1-AF647 and HLA.A2/CMV-AF488 diffused freely with a diffusion constant *D* ranging between 1.0 and 1.2 µm^2^s^-1^ at 37°C (Figure 1C, Figure S1B, S1C).

In initial studies, we isolated CMVsp-CD8+ central memory T-cells from peripheral blood mononuclear cells (PBMCs) of healthy HLA.A2-positive and CMV-seropositive donors. These were in turn lentivirally transduced with a CAR construct featuring a scF_V_ derived from the ROR1-reactive R12 mAb, an IgG4-Fc hinge region and a CD28 transmembrane domain, which was followed by a cytoplasmic tail featuring a 4-1BB co-stimulatory and a CD3ζ activation domain (Figure S1D) (Hudecek et al., 2013). In selected experiments, T-cells also expressed lentivirally-transduced tyrosine kinase ZAP70, which was genetically fused to GFP (ZAP70-GFP) to serve as a probe for CAR- and TCR-proximal signaling. As a direct consequence of canonical antigen receptor-triggering, cytoplasmic ZAP70 becomes recruited to phosphorylated immunoreceptor tyrosine activation motifs (ITAMs) present within the cytoplasmic tails of the CAR and the TCR-CD3 complex (Alberola-Ila et al., 1997). We hence employed synaptic enrichment of ZAP70-GFP within the evanescent TIR field as quantitative readout for CAR- and TCR-proximal signaling (Bunnell et al., 2002).

Functional integrity of the corresponding cell products was verified via antigen-dependent cytotoxicity, cytokine release and proliferation assays (Figure S1E, F and G). To assess the applicability of the SLB-based platform in a physiologically meaningful manner, T-cells were confronted with SLBs functionalized with ICAM-1, B7-1 and either ROR1 or HLA.A2/CMV. CMVsp-ROR1-CAR-T-cells engaged both ROR1 and HLA.A2/CMV immediately upon making SLB contact, as was indicated by rapid antigen enrichment within their synapses (Figure 1D, 2^nd^ row). Antigen binding coincided with vigorous cell activation as witnessed by the synaptic recruitment of ZAP70-GFP (Figure 1D, 3^rd^ row) and by robust calcium signaling monitored via the ratiometric calcium-sensitive dye Fura-2 (Figure 1D, 6^th^ row). Furthermore, we consistently observed in a strictly antigen-dependent manner the transient synaptic polarization of lytic granules (visualized using LysoTracker Deep Red) towards the evanescent TIR field (Figure 1D, 4^th^ row).

We hence conclude that SLBs functionalized with the appropriate set of proteins promoted antigen engagement as well as intracellular signaling pathways driving synapse formation and attack of the target. Furthermore, the use of SLBs as reconstituted surrogate APCs supported quantitative imaging of antigen recognition by living CAR-T-cells at high spatiotemporal resolution while affording considerable experimental flexibility.

### CAR-T cells require ∼1000 times more antigen than CMVsp-CTLs for activation

Both CD4+ and CD8+ T-cells mobilize calcium, undergo lineage commitment and carry out effector functions in response to fewer than 10 antigenic pMHCs on the surface of APCs (Huang et al., 2013; Irvine et al., 2002; Purbhoo et al., 2004). In contrast, thresholds to activate CAR-T-cells were estimated to vary between 100 and 3000 antigens per target cell (Bluhm et al., 2018; Watanabe et al., 2015). While reduced antigen sensitivity may help avoid cytokine storms, off-tumor targeting and premature CAR-T-cell exhaustion, it may also provide a niche for the emergence of antigen escape tumor variants, which have been recently observed at rising frequencies as a complication of CAR-T-cell therapy.

To quantitate and compare the number of antigen molecules needed to activate CAR-T-cells and CMVsp-T-cells, we confronted CMVsp-ROR1-CAR-T-cells with increasing densities of either ROR1 or HLA.A2/CMV on the SLB. We then recorded for a total of 15 minutes antigen-triggered changes in intracellular calcium, which precedes most T-cell effector functions and is a prerequisite for activation-induced changes in gene transcription, cytotoxicity and cytokine release (Pores-Fernando and Zweifach, 2009). As shown for two representative cells encountering their nominal antigen (ROR1 or HLA.A2/CMV, Figure 2A**, upper panel**), cytoplasmic calcium levels peaked in T-cells immediately after making first SLB contact and then reached a plateau (Figure 2A**, lower panel**). For automated high throughput analysis, we applied a python-based scoring algorithm developed in-house to track all cells and to quantitate the percentage of activated cells, i.e. those exhibiting a Fura-2 ratio equal to or above 1.5 for at least 3 minutes (for more details refer to the Methods section). As shown in Figure 2B and **C**, the minimum antigen density required for activation was 0.03 molecules µm^-2^ for HLA.A2/CMV (30% activated cells) and 30 molecules µm^-2^ for ROR1 (12% activated cells). Hence, CMVsp-ROR1-CAR-T-cells needed more than 1000 times more ROR1 molecules than HLA.A2/CMV complexes to mount not only a comparable calcium but also interferon-γ response.

**Figure 2:**
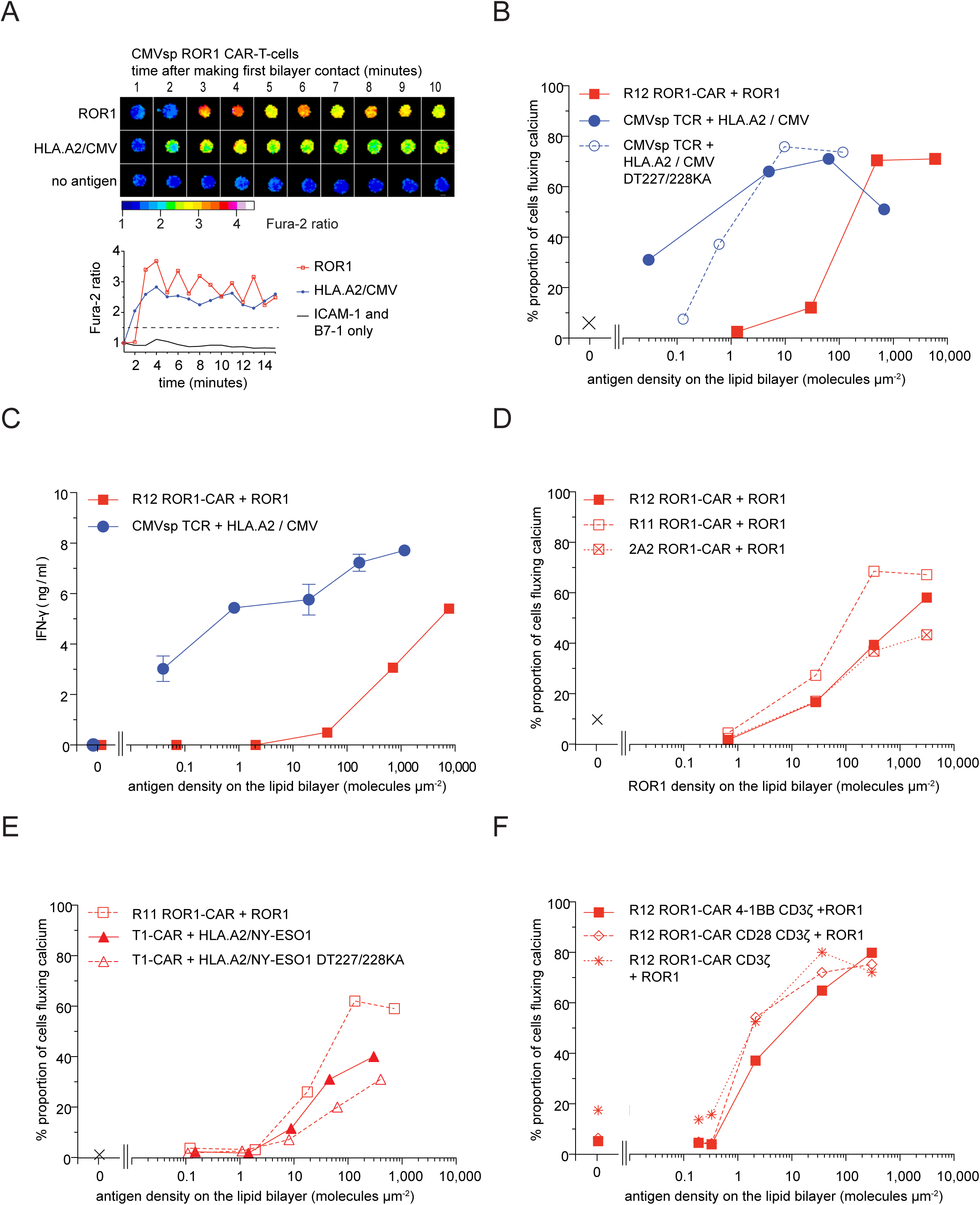
CMVsp-T-cells detect antigen more efficiently than CAR-T-cells by 2-3 orders of magnitude. (**A**, top panel) Fura-2 time-lapse microscopy reveals changes in intracellular calcium concentrations for three representative CMVsp-ROR1-CAR-T-cells confronted with SLBs featuring either ROR1 or HLA.A2/CMV as antigen, or accessory proteins only (scale bar; 5 µm). (**A**, bottom panel) Normalized Fura-2 ratio values are shown for the three cells in the top panel as a function of time. The dashed line indicates the Fura-2 ratio threshold, above which cells were considered activated. (**B**, **D**, **E, F**) Fura-2-loaded CMVsp-ROR1-CAR-T-cells expressing the indicated CAR constructs were confronted with SLBs displaying ROR1 molecules, cognate HLA.A2/peptide complexes or cognate but CD8 binding-deficient (DT227/228KA) HLA.A2/peptide complexes at indicated densities. Cells exhibiting normalized Fura-2 ratios larger than 1.5 for three consecutive minutes were classified as activated. The proportion of calcium-signaling cells was plotted against the corresponding antigen density on the SLB. The first data point (X symbol) denotes a sample of cells confronted with an antigen-free SLB presenting ICAM1 and B7-1. Data from each panel correspond to one donor, total (n=4 donors). Data shown in all panels are representative of 3 or more experiments. The number of cells assayed for each data point ranged from 183 to 3213 (median n=749). (**C**) Quantitation of IFNγ secreted into the media by CMVsp-ROR1-CAR-T-cells interacting with SLBs featuring either ROR1 or HLA.A2/CMV in indicated densities (data points in triplicate; error bars = SD, often too small to be identifiable in the graph).

To investigate underlying causes, we first assessed the role of the CD8 co-receptor, which is known to sensitize CTLs for antigen by binding MHC class I molecules in concert with the TCR (Potter et al., 1987; Purbhoo et al., 2004). Since tumor-associated antigens (TAAs) do not exhibit a CD8 binding site, mechanisms promoting CD8-mediated sensitization are likely absent in CAR-antigen recognition. For a more direct comparison we therefore examined the dose response of CMVsp-ROR1-CAR-T-cells encountering a mutant version of HLA.A2/CMV (DT227/228KA) which no longer binds CD8. As shown in Figure 2B, antigen thresholds required to stimulate CMVsp-ROR1-CAR-T-cells increased in the absence of CD8 engagement from 0.03 to 0.6 molecules µm^-2^, i.e. ∼20-fold. Hence, even when CD8-binding had been abrogated, antigen recognition via CMVsp-TCRs still proved to be 50-times more sensitive than CAR-mediated detection of ROR1. While these observations concur with of rationales involving genetically engineered mechanisms to mimic CD8 function in CAR-T-cells, they also point to the existence of other yet-to-be-identified molecular parameters, which may need to be accounted for when aiming for maximized antigen sensitivity in CAR-T-cell products.

### CAR-T-cell antigen sensitivity is largely unaffected by the epitope and the type of antigen targeted

We next assessed the extent to which the epitope and its position within the antigen influenced detection by CAR-T-cells. To this end we compared the calcium response of T-cells modified with CARs exhibiting scF_V_s derived from the R12 mAb (used earlier in CMVsp-ROR1-CAR-T-cells), the R11 mAb and the 2A2 mAb, which target ROR1 at different epitopes (Figure S2A) (Baskar et al., 2012; Yang et al., 2011). As shown in Figure 2D, dose responses were comparable for the different ROR1-CAR-T-cells, indicating that the position of the epitope relative to the plasma membrane was not decisive for attenuated CAR-signaling output.

Earlier work has shown that length dimensions of TCRs, their nominal pMHC-ligands and accessory proteins within the immunological synapse define optimal kinetic segregation of receptors and kinases from phosphatases for productive signaling (Choudhuri et al., 2005). We verified via immunofluorescence that CAR-antigen micro-clusters were efficiently segregated from CD45, a phosphatase targeting the TCR-CD3 complex, on fixed SLB-stimulated T-cells (Figure S2B). Furthermore, we had previously demonstrated that varying the length of CARs according to the location of the epitope within the antigen had considerable implications for overall CAR performance (Hudecek et al., 2013). However, the extent to which spatial dimensions of the antigen itself play a role in CAR-mediated recognition was unclear.

To find out whether the size of ROR1, which is considerably larger than that of a pMHC molecule, affected CAR-T-cell antigen sensitivity, we substituted the ROR1-specific R11 scF_V_ with a scF_V_ derived from the T1-antibody and binding to HLA-A*0201 complexed with the peptide NY-ESO1 (HLA.A2/NY-ESO1) (Stewart-Jones et al., 2009). As shown in Figure 2E, T-cells expressing the T1 NY-ESO1-CAR or the R11 ROR1-CAR showed similar calcium responses in an antigen density-dependent manner, suggesting that antigen sensitivity was mostly - if not solely - affected by the CAR architecture itself rather than by the type and size of the targeted antigen.

Of note, confronting T1 NY-ESO1-CAR-T-cells with the HLA.A2(DT227/228KA)/NY-ESO1 mutant, which no longer binds CD8, affected the dose response only moderately (Figure 2E), indicating that CD8-HLA.A2 engagement played a rather limited role in CAR-mediated antigen recognition.

### Co-stimulatory signaling modules do not affect antigen thresholds for CAR-T-cell detection

While it is established that costimulatory modules drive both CAR-T-cell survival as well as CAR-T-cell exhaustion *in vivo* (Long et al., 2015; Maus et al., 2014), the extent to which costimulatory signaling motifs affect antigen sensitivity has not been addressed. We hence compared the calcium response of CAR-T-cells, which expressed ROR1-CARs featuring cytoplasmic CD3ζ alone or together with a membrane proximal CD28- or 4-1BB-derived signaling domain (Figure S1D). A small proportion (10-15%) of the T-cells expressing ROR1-CARs lacking a costimulatory module gave rise to spurious calcium signaling without antigen stimulation (Figure 2F). However, sensitivities towards ROR1 were largely comparable among CAR-T-cells featuring CARs with a CD28- or a 4-1BB-derived signaling module, as well as CARs lacking a costimulatory signaling module.

### Synaptic antigen engagement is more pronounced in CAR-T-cells than in CMVsp-T-cells despite moderate CAR surface expression

Because ectopic CAR expression amounted to only 25 to 30% of endogenous TCR expression levels (Figure S3), we assessed whether CAR-antigen binding was functionally limiting when antigen was present at low densities. Since SLB-resident proteins diffuse freely at high speed unless engaged by a receptor expressed on the T-cell side, the number of fluorescently-labeled antigens interacting at any given time with CARs or TCRs can be readily inferred from the magnitude of the fluorescence observed within the synapse (Huppa et al., 2010). Using this rationale, we compared the absolute number of antigens recruited via CARs or TCRs as a function of the density of the antigen present on the SLB prior to T-cell addition (Figure 3A, B). Reflective of their higher binding affinity, ROR1-CAR-T-cells recruited at low antigen densities ∼14 times more antigen than CMVsp-CTLs. Also, as was expected, the number of TCR-engaged pMHCs gradually approached that of CAR-bound ROR1 with increasing antigen densities (Figure 3A, B). Intriguingly, CAR-antigen binding was clearly detectable at antigen densities lower than 10 molecules per µm^2^ yet did not result in any measurable calcium flux (Figure 2B, D, E and F). We therefore conclude that despite lower CAR expression levels synaptic antigen binding was more efficient in CAR-T-cell synapses and *per se* not a limiting factor for CAR-T-cell responsiveness.

**Figure 3:**
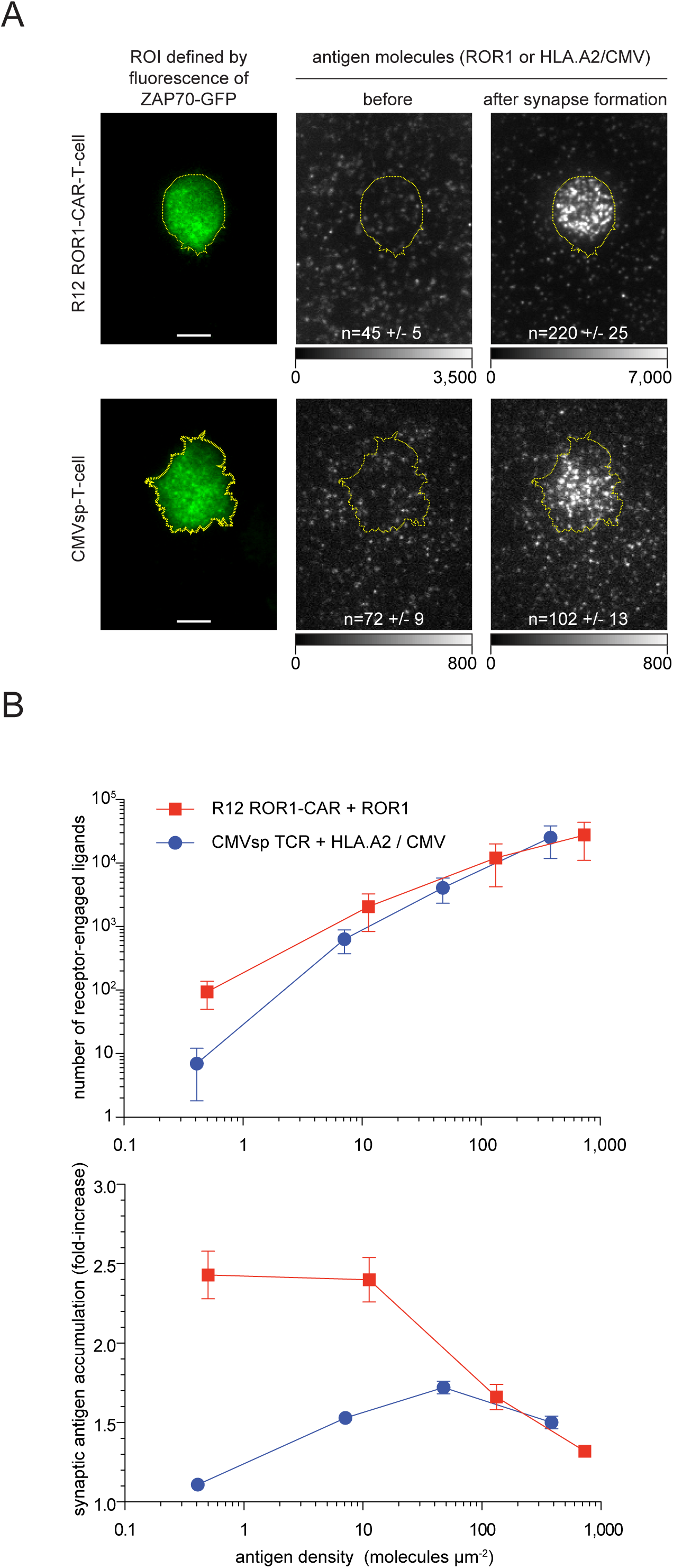
CAR-T-cells engage more antigen than CMVsp-T-cells. (**A**) Synaptic recruitment of fluorophore-labeled antigens was quantitated within a region of interest (ROI) by subtracting the number of fluorescently-labeled SLB-resident antigens prior to synapse formation from the number of antigens after synapse formation. ROIs indicate the synaptic area as defined by the ZAP70-GFP signal recorded in TIR mode (yellow dashed demarcation line). Displayed examples are part of the quantitation shown in (B) (data points referring to lowest antigen densities; +/-indicates SD; scale bar = 5µm). (**B**, upper panel) Absolute numbers of ROR1 and HLA.A2/CMV molecules accumulated within the synapses of ROR1-CAR-Tcells and CMVsp-T-cells were plotted against the density of SLB-resident antigen. Error bars indicate SD. The number of synapses assayed per data point ranged between 24 and 42 (median =32). (**B**, lower panel) The degree of synaptic antigen accumulation was plotted against the density of SLB-resident antigen.

### Synaptic recruitment, activation and release of ZAP70 is compromised

Having ruled out inefficient synaptic antigen engagement as a possible cause for attenuated CAR signaling output, we focused our analysis on signaling events immediately downstream of antigen receptor engagement. ITAM-phosphorylation within the cytoplasmic tails of CARs and TCR-associated CD3 subunits by lymphocyte-specific protein tyrosine kinase (Lck) is the first biochemical change detectable after antigen binding (Chan et al., 1992). To monitor Lck-mediated signaling we quantitated the recruitment of cytoplasmic ZAP70-GFP to phosphorylated ITAMs. For this ROR1-CAR-T-cells and CMVsp-CTLs were sorted by FACS for equivalent ZAP70-GFP expression. FACS-sorted cells were dropped onto SLBs and after acquiring an initial GFP-image, we applied a photo-bleaching pulse in TIR mode to ablate ITAM-associated ZAP70-GFP fluorescence. This was followed by an 800-millisecond recovery phase to determine in a subsequent GFP image the background fluorescence arising from rapidly-diffusing, non-recruited cytoplasmic ZAP70-GFP (for more details refer to Methods section). We simultaneously quantitated the synaptic enrichment of ROR1 and HLA.A2/CMV as a means of correlating the number of ligated antigen receptors to ZAP70 recruitment (Figure 4A). As shown in Figure 4B, significantly more ZAP70-GFP translocated to the synapse at low antigen densities in response to TCR-ligation, even though considerably fewer HLA.A2/CMV molecules had been engaged by TCRs than ROR1 molecules by CARs (Figure 4 A, B).

**Figure 4:**
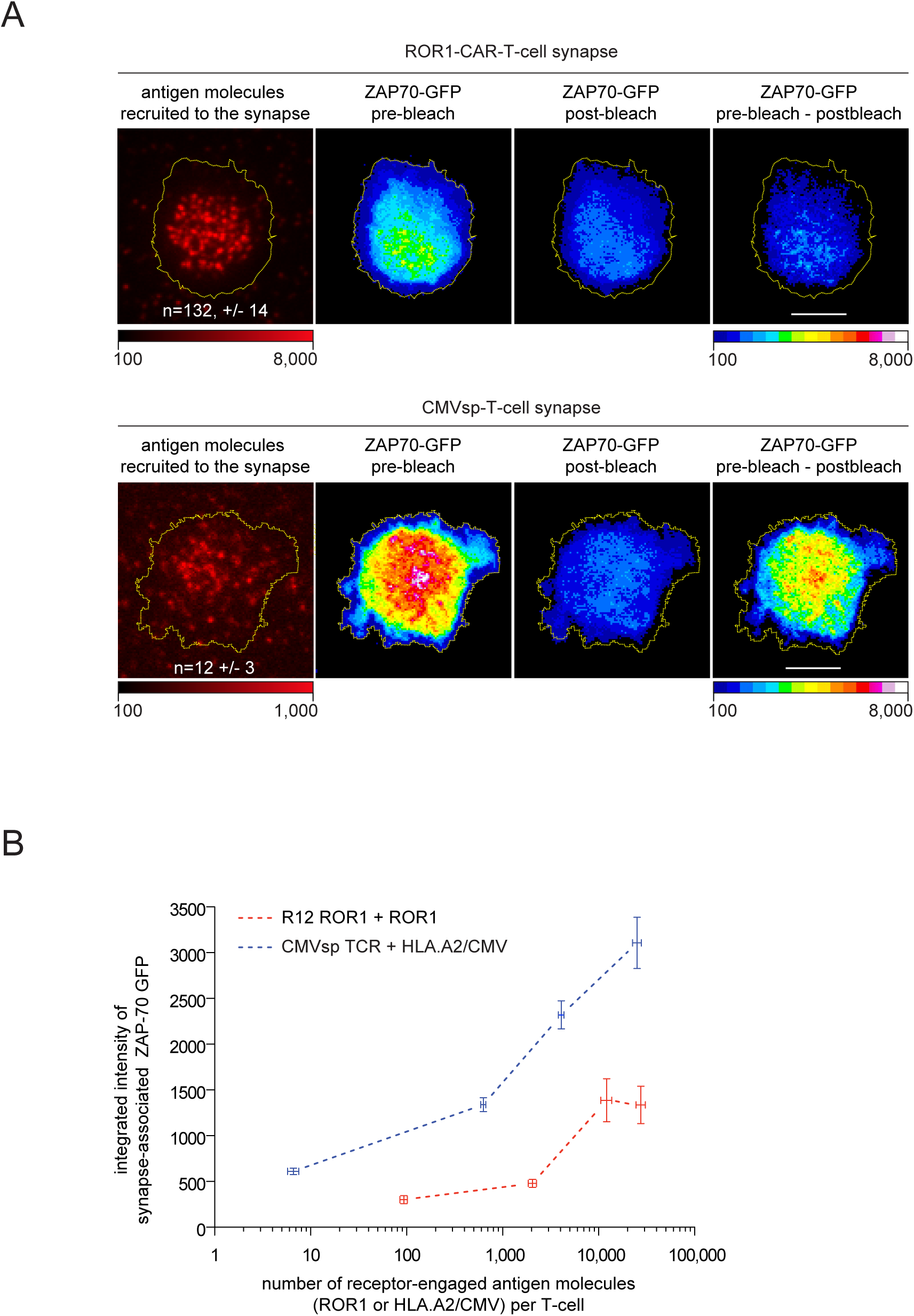
Antigen-ligated CARs recruit ZAP70 less efficiently than antigen-engaged TCRs. **(A)** Representative pseudo-colored TIR images of ZAP70-GFP expressed in a ROR1-CAR-T-cell and a CMVsp-T-cell in contact with antigen-presenting SLBs before photobleaching and 0.8 seconds after photobleaching in TIR mode. The ZAP70-GFP post-bleach images served as reference to determine the pool of cytoplasmic, freely-diffusing ZAP70-GFP, which had not been associated with the synapse, and which was subtracted from the images acquired prior to photobleaching to give rise to intensity values reflective of synapse-associated ZAP70-GFP. ROIs reflective of the immune synapse are indicated by a yellow dashed line. **(B)** Fluorescence intensity values of synapse-associated ZAP70-GFP integrated over the entire synapse were plotted against the number of receptor-engaged ligands per cell at different antigen densities. Error bars indicate SEM. The number of cells per data point ranged from 23 to 42 (median = 29).

Upon ITAM-recruitment ZAP70 becomes activated for further downstream signaling through phosphorylation at positions Y315 and Y319 (Pelosi et al., 1999). To compare the activation state of ZAP70 molecules recruited to either CARs or TCRs, SLB-stimulated T-cells were fixed, permeabilized and stained with a mAb reactive to tyrosine 319-phosphorylated ZAP70 (ZAP70 pY319). Consistent with synaptic ZAP70 activation, enrichment of SLB-resident antigen (ROR1 or HLA.A2/CMV) coincided with ZAP70-GFP recruitment and ZAP70 pY319 staining (Figure 5A). TCR-engagement resulted in a considerably higher degree of ZAP70 pY319 staining, especially at low antigen densities (Figure 5B). To normalize for more proficient ZAP70 recruitment following TCR-engagement, we divided integrated intensity values resulting from ZAP70 pY319 staining by the corresponding synaptic ZAP70-GFP intensity values. As shown in Figure 4B and Figure 5C, antigen**-**ligated CARs recruited not only fewer ZAP70 molecules than pMHC-engaged TCR, but those which had been recruited were also phosphorylated to a lesser extent (0.6-fold). These findings point to ZAP70 activation as another rate-limiting step in CAR-mediated signaling.

**Figure 5:**
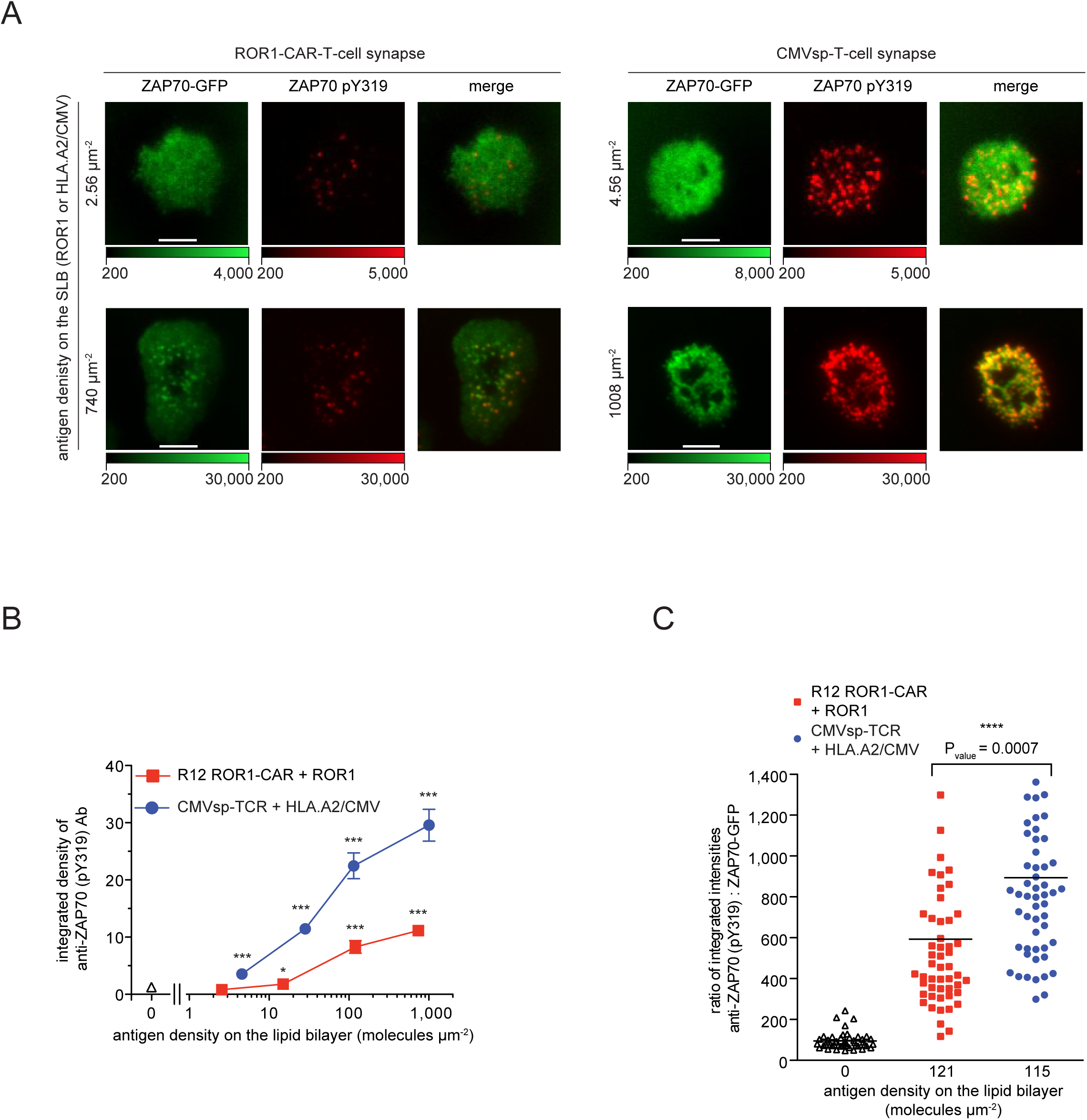
ZAP70 is activated to a lesser extent when recruited to CARs than to TCR/CD3 complexes. **(A)** Accumulation of ZAP70-GFP and ZAP70 phosphorylated at tyrosine 319 (ZAP70 pY319) within synapses of a ROR1-CAR-T-cells and CMVsp-T-cells facing antigen (ROR1 or HLA.A2/CMV) at indicated densities. **(B)** Synapse-associated integrated fluorescence of ZAP70 pY319 was plotted against the density of SLB-resident antigen (ROR1, HLA.A2/CMV). Error bars indicate SEM. The number of cells assayed per data point ranged from 37 to 81 (median = 50). *P* values were calculated by an unpaired two sample Student t-test employing a confidence interval of 95%, comparing the ZAP70 pY319 signal observed in T-cells contacting an antigen-free SLB with that observed in T-cells engaging antigen-presenting SLBs at indicated densities. \**p* < 0.05, \*\**p* ≤ 0.005 and \*\*\**p* ≤ 0.0001. **(C)** Synapse-associated fluorescence intensities of ZAP70 pY319 normalized by the fluorescence intensities of ZAP70-GFP to compare the efficiencies with which ZAP70 was phosphorylated once it had been recruited to the ROR1-CAR-T-cell or the CMVsp-T-cell synapses. Each data point corresponds to one cell and the black bar indicates the geometrical mean. *P* values were calculated as in (B).

When activated, ZAP70 disengages from ITAMs, diffuses laterally within the plasma membrane to target components of the T-cell signalosome and is eventually released into the cytoplasm where it becomes inactivated (Bunnell et al., 2002; Katz et al., 2016). Cycling of ZAP70 is considered integral to signal amplification and a prerequisite for robust calcium fluxing and the initiation of other signaling cascades (Katz et al., 2016). To compare the rates at which ZAP70-GFP molecules became exchanged at CARs and TCRs, we modified a previously published FRAP-based approach (Bunnell et al., 2002). Half of the synaptic area of T-cells, which had been in contact with antigen-loaded SLBs for 5 to 10 minutes, was photobleached in TIR mode with the use of a slit aperture positioned in the excitation beam path (for details please refer to the STAR Methods section). Recovery of synaptic GFP-fluorescence was measured over time to monitor the rate at which bleached receptor-associated ZAP70-GFP was recycled by fluorescent ZAP70-GFP originating from the cytoplasm (Figure 6A). The intensity of the protected area served as an internal control for photobleaching in the course of image acquisition.

**Figure 6:**
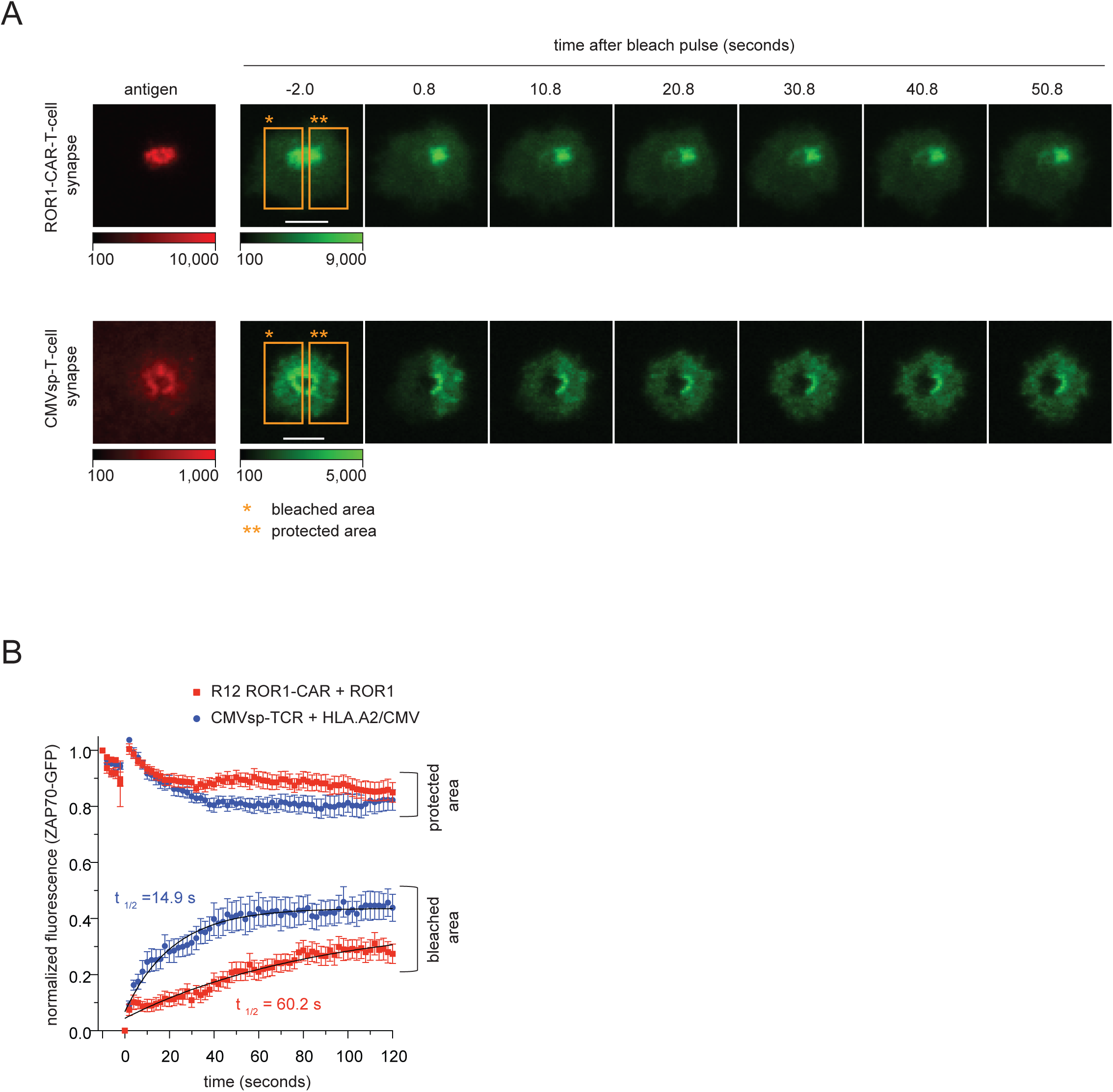
Attenuated dissociation of synapse-associated ZAP70 from CARs. **(A)** Image sequences depicting the recovery of synaptic ZAP70-GFP fluorescence after photobleaching half of a ROR1-CAR-T-cell or CMVsp-T-cell synapse using a slit aperture. Synaptic accumulation of ROR1-AF555 or HLA.A2/CMV-AF555 was used for alignment of the slit aperture prior to photobleaching. Note that only one-half of the synapse was subjected to photobleaching, while the other half was protected during the bleach pulse to serve as internal reference. **(B)** ZAP70-GFP fluorescence intensities of the bleached and protected area of ROR1-CAR-T-cell (n=13) and CMVsp-T-cell synapses (n=16) were normalized against maximal initial ZAP70-GFP fluorescence values and plotted over time following the bleach pulse. Times of half maximal recovery (t_1/2_) were obtained by fitting the normalized ZAP70-GFP signal obtained after photobleaching to an exponential one-phase association model using nonlinear regression. Error bars represent SEM.

As shown in Figure 6B, maximal recovery of GFP-fluorescence was 1.5 times higher in T-cells stimulated via CMVsp-TCRs compared to T-cells stimulated via ROR1-CARs. Furthermore, the rate of recovery was four times faster for TCR-triggered synapses (t_1/2_ of recovery = 15 seconds, n=16) than for CAR-triggered synapses (t_1/2_ of recovery = 60 seconds, n=13), which reflected slower ZAP70-GFP dissociation at the site of activated CARs and consequently less efficient signal amplification at the level of ZAP70. We hence conclude that inefficient Lck-mediated ITAM-phosphorylation and ZAP70-activation limit CAR-T-cell responsiveness to low antigen levels.

## DISCUSSION

The emergence of antigen escape tumor variants with reduced or even complete loss of antigen expression constitutes a severe complication associated with current CAR-T-cell based therapies (Evans et al., 2015; Orlando et al., 2018; Ruella and Maus, 2016). In particular, targeting cancers with highly heterogeneous TAA-expression via state-of-the-art CAR-T-cells remains to this date a challenge (Fry et al., 2018; Perna et al., 2017). While the strategy to “hit hard and early” through the transfer of larger numbers of CAR-modified central memory T-cells may - if applicable - help to increase overall survival rates, it does not provide full protection from outgrowth of tumor variants with low TAA expression.

A central yet unfulfilled need is to endow CAR-T-cells with a tunable capacity to detect TAAs, if necessary, down to single copies per tumor cell, i.e. with a sensitivity towards antigen that matches that of conventional CD4+ and CD8+ T-cells, which carry out effector functions in response to three or even fewer antigenic pMHCs (Huang et al., 2013; Irvine et al., 2002; Purbhoo et al., 2004). This capacity is instrumental for fast pathogen clearance and long-term immunity (Wucherpfennig and Noelle, 2011) and sets a benchmark for the molecular performance of advanced CAR versions, which are devised to eliminate cancer cells displaying one to several hundred TAA molecules (i.e. carrying antigen below the detection level of flow cytometry (Zola, 2004)).

Accelerated CAR development mandates reliable experimental systems affording quantitative, if not single molecule readout in living cells. The SLB-based live-cell imaging platform presented here may very well prove highly relevant in a preclinical setting as it recapitulates cell adhesion, synaptic antigen engagement, antigen receptor-proximal signaling as well as activation-dependent release of cytokines and lytic granules. Testifying to the overall robustness of the assay system, conventional T-cells could readily detect the presence of even a single antigenic pMHC on an SLB, as can be expected for a meaningful readout. Furthermore, we succeeded to our knowledge for the first time in precisely measuring the number of antigens required to activate CAR-T-cells and have identified a major bottleneck in the chain of events triggering CAR-mediated signal transduction.

We found CAR-T-cells superior with regard to synaptic antigen binding, which was anticipated given the high CAR affinity towards antigen. This advantage was however nullified by inefficient downstream signaling, which was already blunted at the level of CAR-ITAM phosphorylation by Lck. As a consequence, antigen thresholds required for CAR-based detection were 2-3 orders of magnitude higher than those required to activate the very same CAR-T-cell product stimulated via their endogenous TCRs. The extent, to which ineffective coreceptor function attenuated CAR-mediated signal transduction, should be investigated in more depth. Indeed, we did not witness a boost in antigen sensitivity for T1-CAR-T-cells, which target an MHC class I molecule (HLA.A2/NY-ESO1) featuring a binding site for CD8 (Figure 2D). This observation suggests that strategies involving coreceptor CARs acting like CD8 to sensitize CAR-T-cells for antigen may require finetuning with regard to the binding affinities and the geometry of antigen engagement by antigen- and coreceptor CARs.

The obvious lack of serial receptor engagement by a single antigen due to the considerably more stable interactions between CARs and their TAAs has been suggested to dampen CAR-T-cell responsiveness (Srivastava and Riddell, 2015). However, previous CAR-T-cell studies have not yet established a role for serial CAR triggering as CARs with lower affinities were reported to diminish rather than enhance T-cell responsiveness (Hudecek et al., 2013; Oren et al., 2014). Our findings provide a novel perspective: benefits arising from serial antigen receptor engagement may rely on the exquisite molecular responsiveness of the TCR-CD3 complex to low affinity ligands, which is not maintained - as we have demonstrated - by state-of-the-art CARs. The penalty to be paid for inefficient CAR signaling may therefore be two-fold: each antigen-receptor interaction does not only give rise to reduced membrane proximal signaling but must also surmount a threshold in binding stability, which in turn no longer permits serial receptor engagement. As a consequence, the number of triggered CARs is at maximum equal to the number of available ligands and becomes a functionally limiting factor when antigen is sparse. In contrast, individual antigenic pMHCs can engage multiple TCR-CD3 complexes with each short-lived encounter giving rise to significantly more potent downstream signaling. Consistent with this concept is the observation that T-cells expressing engineered TCRs featuring CAR-like affinities towards nominal pMHCs are no longer capable of sensitized antigen detection (Thomas et al., 2011).

Our study identifies the magnitude of antigen receptor-triggered ITAM- and ZAP70-phosphorylation as a major limitation in CAR signaling. Yet without clear knowledge of the mechanisms, which have evolved to link extracellular receptor-engagement to intracellular ITAM-phosphorylation, we can only speculate about underlying structural deficits in empirically engineered CARs. Of note, we did not observe significant differences in CAR-proximal responsiveness (Figure 2F) between T-cells modified with ROR1-CARs lacking or featuring a costimulatory signaling module. However, any strategy to boost CAR-T-cell antigen sensitivity may be compromised by lining up co-stimulatory (signal 2 and 3) and TCR-like (signal 1) signaling in series as a result of premature signal convergence and also because ITAMs are invariably placed more distally from the plasma membrane with yet unexplored consequences for Lck accessibility.

The nature of the transmembrane domain may furthermore turn out critical in positioning the signaling modules in adequate vicinity to activating enzymes such as Lck. Palmitoylation of Lck at two N-terminal cysteine residues is critical for functional integrity within the plasma membrane (Kabouridis et al., 1997; Zimmermann et al., 2010). It is hence conceivable that palmitoylation determines accessibility of Lck to ITAMs upon ligand engagement in a yet-to-be-defined fashion, which may in turn be affected by the biophysical properties of the CAR transmembrane domain and the makeup of its local lipid environment.

Other important differences between TCR-CD3 complexes and CARs concern the number of ITAMs per antigen-binding unit (TCR-CD3 complex = 10; CAR = 3-6), their diversity (ITAMs derived from CD3γ, δ, ε and ζ for the TCR, ITAMs from ζ for the CAR) and their three-dimensional arrangement towards one another. Of note, upon ligand binding the proline-rich domain in the cytoplasmic tail of TCR-associated CD3ε (Gil et al., 2002), which is not present in state-of-the-art CARs, recruits the adaptor molecule Nck and may affect signalosome assembly. Furthermore, signal amplification through autophosphorylation of ZAP70 (Neumeister et al., 1995) may require specific juxtaposition of ITAM-engaged ZAP70 molecules that may no longer be preserved in CARs. Increasing evidence points to changes within the TCR-CD3 quaternary structure as a trigger for TCR-signaling with no equivalent mechanism for current CAR-based systems (Parrish et al., 2015).

Hence, strategies to improve antigen detection capabilities by CAR-T-cells are likely iterative in nature and may involve (i) the adaptation of CARs to the architectural framework of the TCR-CD3 complex, for example by linking the TAA-specific scF_V_ to either the TCR or one of CD3 subunits and (ii) the separation of signal 1 from signals 2 and 3 through simultaneous expression of two or more CARs engaging the same or different TAAs. Fine-tuning antigen binding kinetics to allow for serial CAR engagement by a single antigen may further enhance such CAR formats. If successful, future developments of sensitized CARs will set high demands on achieving tumor specificity to avoid off-target toxicities.

In summary, highly resolved live-cell imaging allowed us to gauge antigen sensitivities exhibited by CAR-T-cells and to identify inefficient CAR-proximal Lck- and ZAP70-signaling as major shortcomings of current CAR formats. Given its molecular readout and ease of use, we consider the featured SLB-based imaging approach instrumental for rational development, refinement and adjustment of CAR formats with enhanced antigen sensitivity, tumor specificity and other properties deemed important for effective, safe and personalized treatment of malignancies.

## CONTRIBUTIONS

VG, JR, MH and JH conceived the project and wrote the manuscript. VG conducted and analyzed all imaging-related and biochemical experiments. JR generated all T-cell lines and conducted classical immunological assays. IP wrote analysis software. SK carried out initial experiments and developed the CAR-T-cell-specific SLB system. LS produced HLA.A2 and HLA.A2(DT227/228KA) imaging probes. HE and HS contributed important ideas.

## ACKNOWLEDGEMENTS

This work was supported by the Austrian Science Fund (FWF) through the P 25775-B2 project (VG, SK, LS and JH), the Vienna Science and Technology Fund (WWTF) LS14-0301(VG, JH) and by grants from the German Cancer Aid (Deutsche Krebshilfe e.V.) through the Max Eder Program 110313 (MH). We thank René Platzer (Medical University of Vienna) for help regarding single molecule tracking, Omer Dushek (Oxford University) for sending us the T1-CAR-constructs and Ulrich Jäger and Nina Worel (both Medical University of Vienna), Manfred Lehner (St. Anna Children’s Cancer Research, Vienna) and Armin Rehm (Max Delbrück Center for Molecular Medicine, Berlin) for constructive criticism.

## SUPPLEMENATAL FIGURE LEGENDS

**Figure S1:**
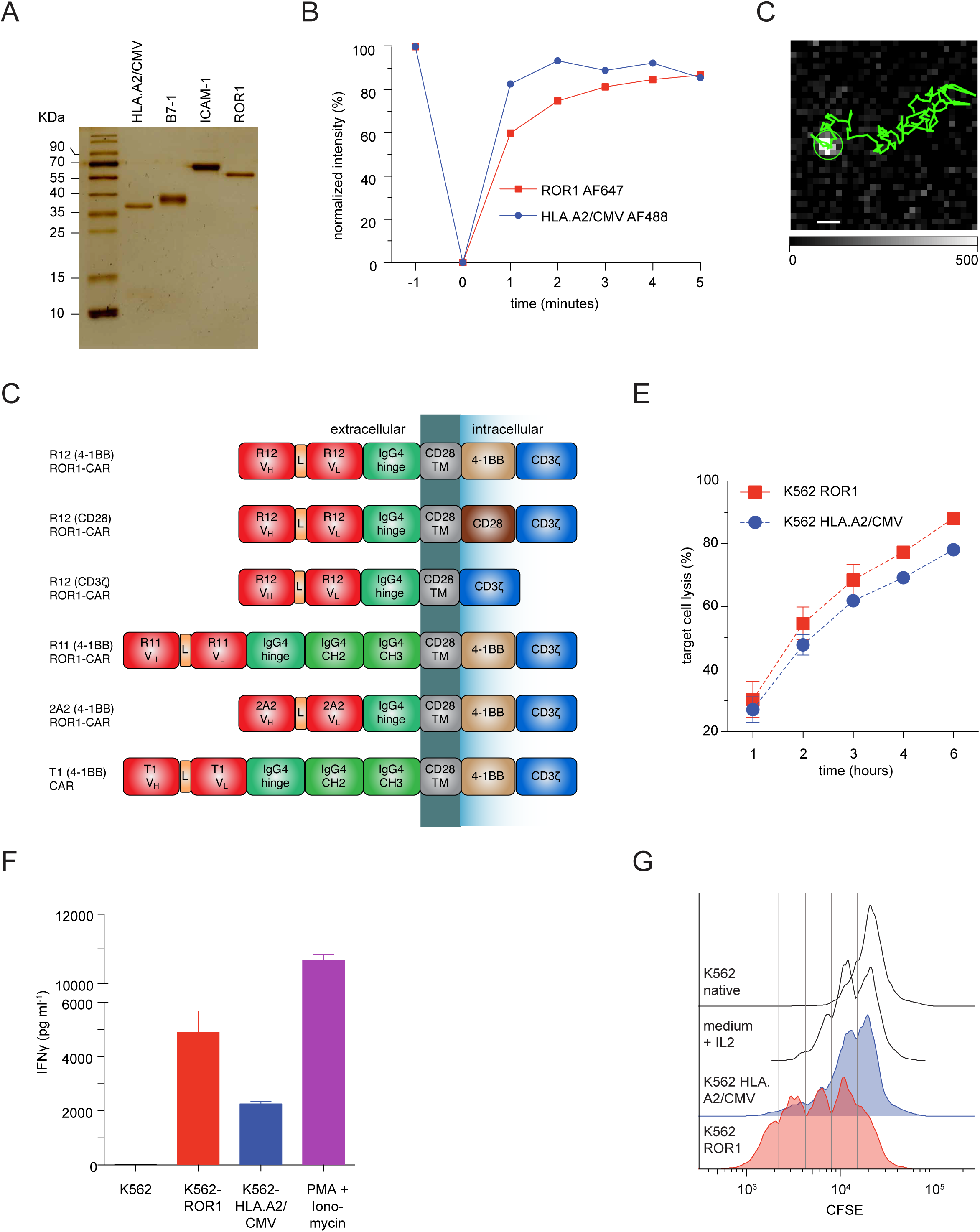
Biochemical and imaging-based analysis of SLB-embedded proteins as well as *in vitro* analysis of cytotoxicity, IFNγ secretion and proliferation of CMVsp- and ROR1-CAR-T-cells. **(A)** 12.5% SDS PAGE analysis of the recombinant proteins employed for SLB functionalization. **(B)** The immobile fraction of SLB-anchored proteins was determined via <u>F</u>luorescence <u>R</u>ecovery <u>A</u>fter <u>P</u>hotobleaching (FRAP). Fluorescence intensities were normalized with regard to initial intensity values and plotted versus time. **(C)** The diffraction-limited fluorescence signal of a single ROR1-AF555 molecule diffusing laterally in an SLB is marked with a green circle. The multi-step single molecule time trajectory is indicated as a green trace (scale bar: 1µm). **(D)** Schematic outline of CAR-constructs employed in this study. **(E)** Cytotoxicity of CMVsp-ROR1-CAR-T-cells co-cultured with K562-ROR1 or K562-HLA.A2/CMV target cells (10:1 E:T ratio) over time. Specific lysis was determined using a bioluminescence-based assay (Wang et al., 2011). Error bars indicate SD. **(F)** IFNγ secretion of CMVsp-ROR1-CAR-T-cells co-cultured for 24 hours with K562-ROR1 or K562-HLA.A2/CMV target cells (4:1 ratio) or in the presence of PMA and ionomycin. IFNγ concentration in the supernatant was analyzed by ELISA. Error bars indicate SD. (G) Proliferation of CFSE-labeled CMVsp-ROR1-CAR-T-cells co-cultured for 72 hours with K562-ROR1 or K562-HLA.A2/CMV target cells (4:1 ratio) or high IL-2 concentrations. As indicated, the decrease in fluorescence was reflective of the number of cell divisions.

**Figure S2:**
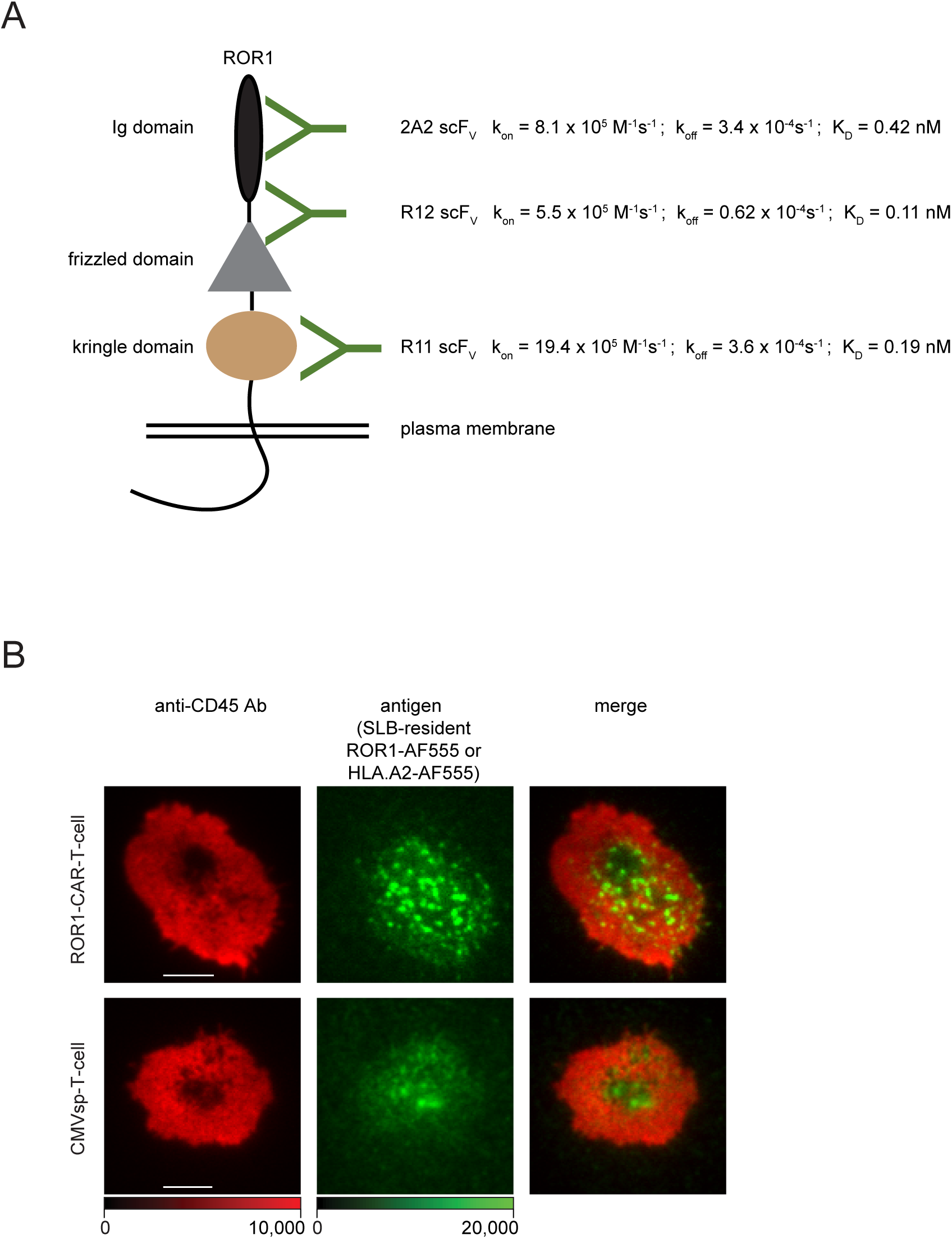
Epitopes targeted by ROR1-specific CARs and CD45 segregation patterns observed in immunological synapses of R12 ROR1-CAR-T-cells and CMVsp-T-cells. **(A)** Schematic representation of ROR1-epitopes targeted by R12, R11 and 2A2 CARs with previously published binding constants (Baskar et al., 2012; Yang et al., 2011). **(B)** Representative images of immunological synapses of an R12-ROR1-CAR-T-cell and a CMVsp-T-cell confronted with SLB-embedded antigen as indicated. Note the segregation of CD45 from synaptic areas in which antigen binding is prominent.

**Supplemental Figure 3.**
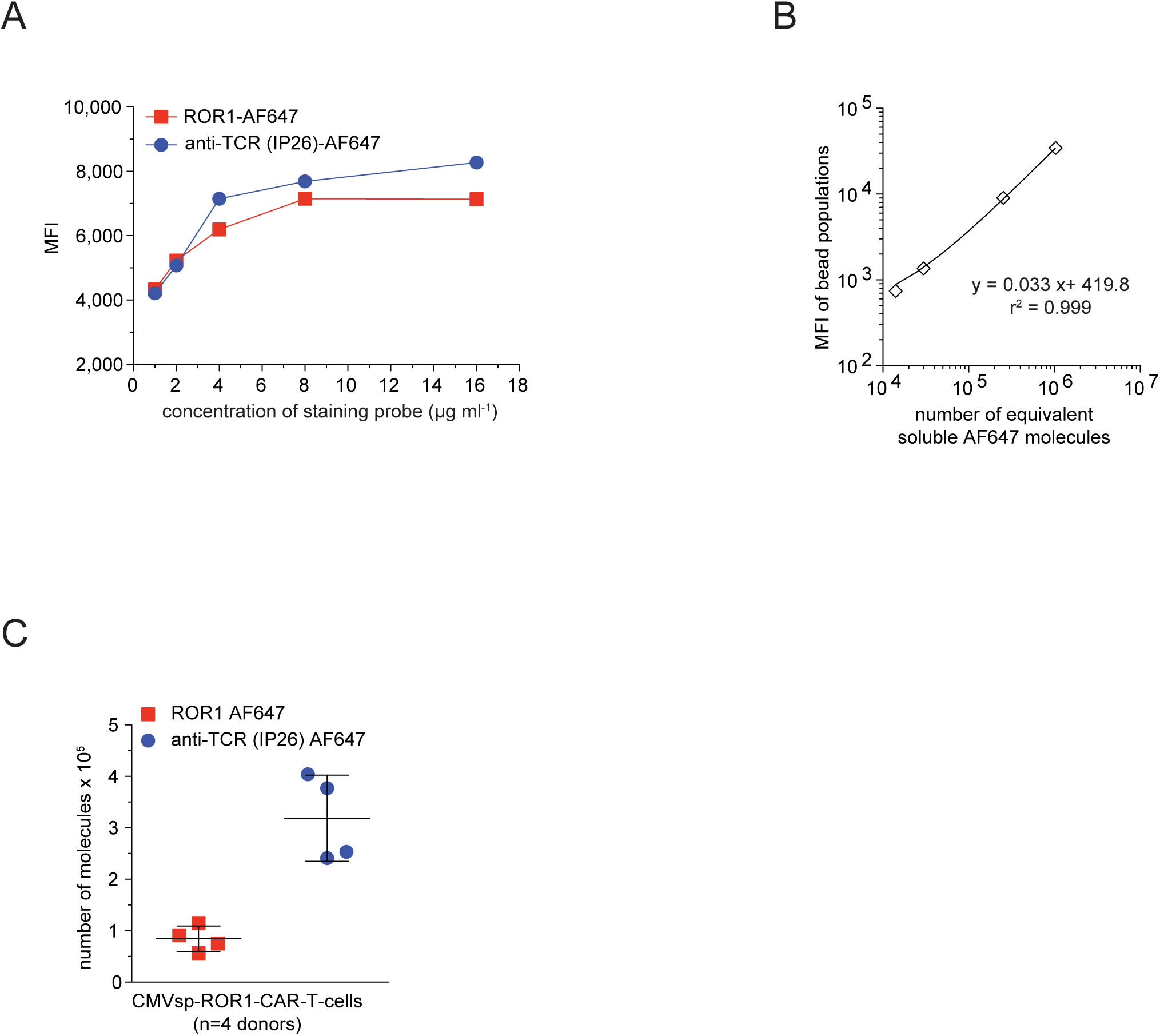
Determination of TCR and CAR surface densities. **(A)** CMVsp-ROR1-CAR-T-cells were labeled with either recombinant ROR-AF647 or AF647-conjugated anti-TCR mAb (clone IP26) to determine concentrations required for label saturation. **(B)** Flow cytometric calibration of AF647 intensity values with the use of Quantum™ Alexa Fluor^®^ 647 MESF beads. Resulting MFIs were plotted against the number of equivalent AF647 fluorophores. **(C)** Quantitation of CARs or TCRs on the surface of CMVsp-ROR1-CAR-T-cells. Error bars represent SD.

## STAR METHODS (Gudipati et al.)

### Constructs for lentiviral gene transfer

ROR1-specific scFvs were generated by genetically fusing the V_H_ and V_L_ domains of antibodies R12, R11 and 2A2 via a (G_4_S)_3_ linker (Baskar et al., 2012; Yang et al., 2011); scFvs were cloned into the lentiviral vector epHIV7 upstream of a short or long IgG4-Fc-derived hinge domain, a CD28 transmembrane domain, and intracellular signaling domains (4-1BB, CD28, CD3ζ). The construct encoding NY-ESO-1-specific T1 scFv (Stewart-Jones et al., 2009) was a kind gift from Dr. Omer Dushek (Oxford University) and was cloned into the CAR framework with a long IgG4-Fc-derived hinge domain, a CD28 transmembrane domain, and 4-1BB and CD3ζ signaling domains. All CAR-encoding expression vectors contained an EGFRt (EGFR truncated) transduction marker separated by a viral T2A cleavage sequence (Wang et al., 2011). The ZAP70-GFP fusion construct included the complete coding sequence of human ZAP70 (UniProt: P43403) which was connected via a GSG linker to monomeric GFP (A206K mutation) (Zacharias et al., 2002) and cloned into the lentiviral vector epHIV7 upstream of a viral T2A cleavage sequence and HER2t (HER2 truncated, UniProt: P04626, amino acids 1-652) transduction marker.

### Generation of CMVsp-T-cells and CAR-T-cells

CD8^+^ memory T-cells were isolated by magnetic cell separation (MACS® MicroBeads Technology, Miltenyi) from PBMCs of healthy donors or from PBMCs of HLA.A2, CMV serum positive donors expressing a TCR recognizing the CMV pp65-derived peptide NLVPMVATV. Anti-CD3/CD28 Dynabeads® (Thermo Fisher Scientific) were used for polyclonal T-cell stimulation, while CMVsp-T-cells were stimulated at a ratio of 3:1 with autologous PBMCs irradiated at 30 Gy and pulsed with PepTivator® CMV pp65 peptide pool (Miltenyi). T-cells were cultivated at 37°C and 5% atmospheric CO_2_ in RPMI 1640 medium supplemented with 25 mM HEPES, 10% human serum, 100 U/ml penicillin/streptomycin, 50 µM 2-mercaptoethanol (Thermo Fisher Scientific) and 50 IU/ml recombinant IL-2 (Novartis). On the following day, polybrene (Sigma) was added to the media at 5 µg/ml and T-cells were transduced with lentiviral particles to stably integrate ZAP70-GFP or CAR-encoding genes. The anti-CD3/CD28 Dynabeads® were removed on day 7 post transduction and the percentages of CAR-, ZAP70-GFP- and CMVsp-TCR-positive cells were determined by staining with ROR1-AF647, anti-HER2-AF647 antibody and HLA.A2/CMV-strep plus Strep-Tactin-APC (IBA-Lifesciences, Germany), respectively. Transgene-positive T-cells were enriched via the transduction markers EGFRt and HER2t using biotin-labeled anti-EGFR and anti-HER2 antibodies, respectively, and anti-biotin MicroBeads (Miltenyi). For expansion, CMVsp-T-cells and T1-CAR-T-cells were co-cultured with K562-HLA.A2 cells irradiated at 80 Gy and pulsed with CMV pp65 peptide or NY-ESO-1 peptide, while ROR1-CAR-T-cells were co-cultured with irradiated K562 cells, which had been transduced to express the extracellular domain of ROR1 (K562-ROR1) (Hudecek et al., 2010) at a ratio of 4:1. Every second day, half of the medium was exchanged with fresh T-cell medium containing recombinant IL-2 (100 IU/ml) (Novartis), and experiments were conducted after 8 to 11 days of expansion and 12 or more hours after medium exchange.

### Functional *in vitro* analysis of T-cells

Cytolytic activity of T-cells was analyzed against K562 cells expressing firefly luciferase (Brown et al., 2005). Briefly, effector cells (50×10^3^) were mixed with target cells (5×10^3^) and D-luciferin substrate was added to measure the luminescence after 1 to 6 hours. IFNγ secretion was quantitated through ELISA (BioLegend) of supernatants from T-cell cultures co-incubated for 20 hours with K562 target cells at a ratio of 4:1. Proliferation of CFSE-labeled T-cells was assessed after 72-hours co-culture with K562 target cells at a ratio of 4:1 by flow cytometry.

### Molecular cloning, protein expression and purification

cDNA encoding the extracellular domains excluding the leader sequence of the human proteins ROR1 (UniProt: Q01973), B7-1 (UniProt: P33681) and ICAM-1 (UniProt: P05362) was amplified by PCR and inserted into vector pAcGP67 (BaculoGold™ Baculovirus Expression System, BD Biosciences). This vector had been modified so that all proteins carried a c-terminal AVI-tag followed by a 3C protease cleavage site and a poly-histidine-tag harboring 12 consecutive histidine residues (H_12_). High Five™ cells (BTI-TN-5B1-4, Thermo Fisher Scientific) were infected with viral particles produced with the Baculovirus Expression System according to the manufacturer’s instructions (BD Biosciences). The supernatant was harvested 3 to 4 days after infection at 90% cell viability by centrifugation and filtration (0.45 µm) and dialyzed against PBS by tangential flow filtration (Minimate™ TFF System equipped with 10 kDa T-Series cassettes, Pall Corporation). For buffer exchange, supernatant containing the expressed protein was first concentrated 8-fold and then diluted 8-fold with PBS, which was repeated twice. The buffer-exchanged supernatant was concentrated 8-fold one more time and then subjected to Ni^2+^-NTA agarose chromatography (HisTrap excel, 2 x 5 ml columns connected in series, GE Healthcare). Protein was eluted with PBS/300 mM imidazole (pH 7.4) and subjected to size exclusion chromatography (Superdex 200 10/300 GL, GE Healthcare) and anion-exchange chromatography (Mono Q 5/50 GL, GE Healthcare). All purification steps were performed on an ÄKTA™ pure chromatography system (GE Healthcare). Eluted fractions were analyzed by SDS-PAGE followed by silver staining. Purified protein was labeled for microscopy if required (see below), adjusted to 50% glycerol/PBS, and snap-frozen in liquid nitrogen for storage at -80 °C until further use.

cDNA encoding the extracellular domains of HLA.A2 (UniProt: P01892) and beta-2-microglobulin (UniProt: P61769) was cloned without the leader sequence into pET-28b and pHN1 vectors, respectively, for expression in *E. coli*. pET-28b was modified to provide a c-terminal H_12_ tag for HLA.A2. HLA.A2-H_12_ and beta-2-microglobulin were expressed as inclusion bodies and refolded in the presence of peptide (i.e. human cytomegalovirus structural protein pp65 amino acid residues 157–165, NLVPMVATV; human cancer testis antigen NY-ESO-1 amino acid residues 157-165, SLLMWITQC) to give rise to HLA.A2/peptide complexes (Clements et al., 2002; Garboczi et al., 1992). After completion of the refolding reaction the protein solution (200 ml) was dialyzed three times against 10 l PBS. Properly conformed pMHC molecules were purified by Ni^2+^-NTA agarose chromatography (HisTrap excel, 2 x 5 ml columns connected in series, GE Healthcare) followed by size exclusion chromatography (Superdex 200 10/300 GL, GE Healthcare). Fractions containing correctly formed HLA.A2/peptide complexes were identified by SDS-PAGE followed by silver staining. Protein used for microscopy was labeled immediately after purification (see below), adjusted to 50% glycerol/PBS, and snap-frozen in liquid nitrogen for storage at -80 °C until further use.

### Fluorophore conjugation

100 µg of protein in PBS was concentrated to 1-2 mg/ml with the use of Amicon® Ultra Centrifugal Filters (Merck), adjusted to pH 8.3 by addition of freshly-prepared NaHCO_3_ (0.1 M final concentration), an^d^ incubated for 13 minutes at 30 °C with NHS ester derivatives of Alexa Fluor 488, Alexa Fluor 555 or Alexa Fluor 647 (all Thermo Fisher Scientific) present in 6-fold molar excess. The protein was subjected to size exclusion chromatography (Superdex 200 10/300 GL, GE Healthcare) to separate protein:dye conjugates from unreacted dye. The degree of labeling was determined by photospectrometry and labeled protein was adjusted to 50% glycerol/PBS, snap-frozen in liquid nitrogen for storage at -80 °C until further use.

### Preparation of glass supported lipid bilayers (SLBs)

SLBs were prepared as described by (Axmann et al., 2015). In brief, 1,2-dioleoyl-sn-glycero-3-[N(5-amino-1-carboxypentyl)iminodiacetic acid) succinyl] [nickelsalt] (Ni-DOGS-NTA) and 1-palmitoyl-2-oleoyl-sn-glycero-3-phosphocholine (POPC) (both from Avanti Polar Lipids) were dissolved in chloroform and mixed in a 1:9 molar ratio. After drying under vacuum in a desiccator overnight, the lipids were re-suspended in 10 ml degassed PBS and sonicated under nitrogen at 120–170 W in a water bath sonicator (Q700, QSonica) for up to 90 minutes until the suspension had lost most of its turbidity. To pellet non-unilamellar vesicles, the suspension was centrifuged for 4 hours at 37,000 rpm and 25 °C using a Sorvall RC M150GX ultracentrifuge using a S150AT-0121 rotor (Thermo Fisher Scientific). The clear supernatant was centrifuged again for 8 hours at 43,000 rpm and 4 °C employing the same centrifuge and rotor. The second supernatant was filtered once through a 0.2 µm syringe filter (Filtropur S 0.2, Sarstedt) and stored under nitrogen at 4 °C for up to 6 months.

Glass slides (24 mm x 50 mm #1 borosilicate, VWR) were immersed in a 2:1 mixture (v/v) of concentrated sulfuric acid and 30% hydrogen peroxide (both Merck) for 60 minutes, rinsed with deionized water and air-dried. Cleaned slides were attached to the bottom of an 8-well Lab-Tek™ chamber (Nunc) with picodent twinsil extrahart (Picodent) until the glue had solidified. The lipid vesicle suspension was diluted 1:20 in PBS and filtered through a 0.2 µm filter. Of these, 200 µl were added per well to form a contiguous SLB within 5 minutes. Chambers were washed twice with 15 ml PBS to remove excessive vesicles. H_12_-tagged proteins of interest were added to the SLBs and incubated for 60 minutes in the dark at room temperature. Finally, chambers were rinsed twice with 15 ml PBS to remove unbound protein.

### Microscopy setup

Microscopy was conducted with two inverted setups, which had been custom-built to allow for calcium and TIR-based imaging. Unless specified otherwise, excitation light was provided by diode lasers featuring 488 nm, 647 nm (both iBeam smart Toptica) and 532 nm (OBIS) laser lines on both microscopy setups, which were also equipped with a custom Notch filter (Chroma Technology) to block reflected stray light of 488 nm, 532 nm or 640 nm from reaching the camera.

i. An inverted microscope (DMI 4000, Leica) was equipped with a chromatically corrected 100X TIR objective (HC PL APO 100x/1.47 OIL CORR TIRF, Leica), a 20X objective (HC PL FLUOTAR 20X/0.50 PH2 ∞/0.17/D, Leica) and in addition to the lasers specified above with a mercury lamp (EL6000, Leica) for Fura-2-based calcium recordings. Furthermore, this microscope included two fast-filter wheels containing 340/26 and 387/11 excitation bandpass filters (both Leica), and ET525/36, ET605/52 and ET705/72 emission bandpass filters present in the emission pathway (all Leica), as well as a motorized multichroic mirror turret equipped with FU2 (Leica) and ZT405/488/532/647rpc beam splitters (AHF).
ii. A second inverted microscope (Eclipse Ti-E, Nikon) was equipped with a chromatically corrected 100X TIR objective (CFI SR Apo TIR 100X Oil NA:1.49, Nikon), a 20X objective (CFI S Fluor 20X NA:0.75, Nikon) and, in addition to specified lasers, a xenon lamp (Lambda LS, Sutter Instrument) providing illumination for Fura-2-based calcium measurements. This microscope featured also a Lambda 10-3 excitation filter wheel (Sutter Instrument) equipped with 340/26 and 387/11 bandpass filters (both AHF), and a Lambda 10-3 emission filter wheel featuring bandpass filters ET510/20, ET605/52, ET700/75 (all AHF). In addition, two polychroic filter turrets allowed for the implementation of two independent beam paths for laser- and lamp-mediated illumination involving zt405/488/532/640rpc and 409/493/573/652 beam splitters (both AHF).

Data were recorded using an iXon Ultra 897 EM-CCD camera (Andor). An 8 channel DAQ-board PCI-DDA08/16 (National Instruments) and the microscopy automation and image analysis software Metamorph (Molecular Devices) were used to program and apply timing protocols and control all hardware components. Data analysis was performed with Fiji image processing package based on ImageJ (Schindelin et al., 2012; Schindelin et al., 2015). Prism 5 (GraphPad) was used for statistical data analysis and plotting graphs.

### Measurements of antigen densities

FRAP analysis and antigen density measurements were performed exactly as described by us previously in a methods paper (Axmann et al., 2015). In brief, within a central ROI of 60×60 pixels and featuring maximal illumination (placed within the central area of the laser illumination spot) a 7×7 pixel ROI was placed around a diffraction limited fluorescent event and a neighboring area free of fluorescence and intensity counts were integrated. Background-corrected intensity values of 50-100 diffraction-limited single molecule fluorescence events were averaged to determine the single-molecule-signal (SMS) at the excitation power density given for single molecule detection. For quantitation of antigen densities on SLBs, illumination power density was reduced by a factor of 10, 100 or 1000 and fluorescence counts were integrated within the central 60×60 pixel ROI, background-subtracted (background had been measured under identical imaging conditions on antigen-free SLBs), divided by the determined single molecule signal and multiplied by the factor, by which the illumination power density had been reduced (e.g. 10, 100 or 1000 by placing an neutral density filter of OD1, OD2 or OD3 in the excitation beam path) to arrive at the number of molecules present within the central ROI, which was divided by the 60 x60 pixel area (1 pixel ≙ 0.0256 µm^2^, 60 x 60 pixels = 3600 pixels ≙ 92.16 µm^2^) to finally arrive at the antigen density.

### Determination of the immobile fraction by FRAP

Fluorescence recovery after photobleaching (FRAP)-based experiments were conducted as published (Axmann et al., 2015) to determine the fraction of immobile SLB-resident molecules. In brief, a circular aperture was placed in the excitation beam path. After recording 2 to 5 images, the SLB was subjected to an intense bleach pulse, which was followed by 5 to 10 images acquired at 1-minute time intervals. Intensity values of the acquired images were normalized against intensities measured in images acquired prior to bleaching.

### Single dye tracing experiments

Diffusion coefficients of labeled SLB-anchored protein were measured as described below. We recorded single molecule trajectories for up to 1000 frames with a 10 milliseconds exposure time and a total time lag of 10.3 milliseconds. Fluorophore localizations were combined into trajectories according to previously published algorithms describing a more accurate method of ascertaining fluorophore sub-pixel localization (Gao and Kilfoil, 2009; Wieser et al., 2007). We calculated the mean square displacement (MSD) describing the average of the square displacements between two points of the trajectory according to

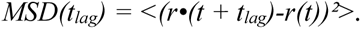

The first three MSD values as a function of t_lag_ were used to calculate the diffusion coefficient *D* for each trajectory by fitting

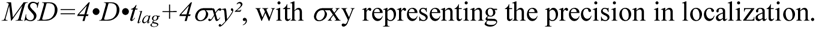

### Cell labeling with LysoTracker™ Deep Red and Fura-2-AM

1×10^6^ cells were incubated in 1 ml conditioned medium supplemented with 50 nM LysoTracker™ Deep Red and 5 µM Fura-2-AM (both Thermo Fisher Scientific) for 30 minutes at 37 °C. The cells were pelleted at 500g and washed twice in 1 ml ice-cold imaging buffer, which consisted of Hank’s Buffered Salt Solution (HBSS) supplemented with 2 mM CaCl_2_ 2 mM MgCl_2_, 2% FCS and 10 mM HEPES (Thermo Fisher Scientific), resuspended in 50 µl imaging buffer and stored on ice for up to 2 hours.

### Monitoring synaptic recruitment of fluorescent antigen, lytic granules and ZAP70-GFP

Images were recorded in TIR-mode using the chromatically corrected 100X objective with high numerical aperture (CFI SR Apo TIR 100X Oil NA:1.49, Nikon). Two to ten minutes after seeding, T-cells were imaged to monitor (i) the redistribution of SLB-resident antigen linked to fluorophore AF555 (using 532 nm laser illumination) and (ii) the recruitment of lytic granules (using 640 nm laser illumination) and ZAP70-GFP (using 488 nm laser illumination) appearing in the TIR-sensitive field of evanescent light.

### Calcium imaging

Changes in intracellular calcium levels were measured via the ratiometric calcium-sensitive dye Fura-2-AM (Thermo Fisher Scientific) as described (Roe et al., 1990). 1×10^6^ T-cells were incubated in 1 ml conditioned medium supplemented with 5 µM Fura-2-AM for 30 minutes at room temperature, washed twice in 1 ml ice-cold imaging buffer (HBSS supplemented with 2 mM CaCl_2_, 2 mM MgCl_2_, 2% FCS and 10 mM HEPES), resuspended in 50 µl imaging buffer and stored on ice for up to 2 hours. For imaging, T-cells were placed in close proximity to the SLB. As soon as the first cells touched the SLB, 510/80 nm emission was recorded with 340 and 387 nm excitation every minute for 15 minutes, visiting 5 different XY-stage positions and using a UV-transmissive 20X objective (CFI S Fluor 20X NA:0.75, Nikon) or (HC PL FLUOTAR 20X/0.50 PH2 ∞/0.17/D, Leica).

For analysis, 340 nm and 387 nm images were subjected to rolling ball background subtraction (Fiji, built-in plugin). Resulting 340 nm stacks were divided by the corresponding 387 nm stacks to generate ratiometric stacks. Individual cells were detected in the 340 nm channel by using Difference of Gaussians detector (DoG detector (Lowe, 2004), tracked in XY and time by using Linear Assignment Problem tracking (LAP tracker (Jaqaman et al., 2008) with the use of the Fiji Trackmate plug-in (Tinevez et al., 2017). XY-plane positions of the detected cells and their corresponding tracks were then applied to ratiometric stack and integrated fluorescence intensity values pertaining to all tracks were recorded. Unless indicated otherwise, parameters (e.g. radius of cell, length and quality of track) were chosen heuristically. Data corresponding to tracked cells were analyzed applying a custom-written python-3 code and python packages (numpy, matplolib, pandas). Intensity values corresponding to individual tracks were smoothened with a Gaussian rolling window filter, whereupon we identified a frame in which the change in intensity was maximal. Upon identifying this frame of activation, the raw data corresponding to the entire time series were normalized to an average intensity value of all frames prior to the frame of activation. After normalization, tracks were classified as activating when the average intensity value of three consecutive frames (i.e. spanning 3 minutes) after the frame of activation was >1.5. The results of each individual track were then combined to characterize the population.

### Quantitation of IFNγ secretion after antigenic stimulation of T-cells on SLBs

Cells (3×10^5^) resuspended in 100 µl imaging buffer were seeded onto to SLBs and incubated at 37 °C for 15 minutes. Following the incubation, 450 µl of RPMI 1640 medium supplemented with 25 mM HEPES, 10% FCS, 100 U/ml penicillin/streptomycin, 2mM L-glutamine and 50 µM 2-mercaptoethanol (Thermo Fisher Scientific) was added followed by a 72 h incubation. The supernatant was than collected and diluted 1:1 with fresh medium and stored as 100 µl aliquots and stored at -80 °C until further use. The amount of IFNγ secreted was then measured by performing an ELISA using a commercially available kit (BioLegend).

### Quantitation of receptor engaged antigens

Images were recorded in TIR-mode with the use of a chromatically corrected 100X objective (CFI SR Apo TIR 100X Oil NA:1.49, Nikon). Prior to cell seeding, a number of 532 nm images of SLB-resident AF555-conjugated antigen was acquired. Cells were then seeded onto the SLB and incubated for 2 minutes. Within the following 8 minutes, up to 20 individual synapses were recorded with 532 nm excitation to quantitate synaptic antigen enrichment. Given the high lateral mobility of SLB-embedded antigens, the number of antigens accumulating underneath a given T-cell corresponds to the number of antigen receptors engaged with a ligand, which we calculated by subtracting initial antigen densities measured prior to cell seeding from synaptic antigen densities. Determined values were multiplied by the synaptic area to arrive at the number of antigen receptors engaged by an SLB-resident antigen (e.g. ROR1 or HLA.A2/CMV).

### Quantitation of synapse-recruited ZAP70-GFP

To discriminate freely diffusing ZAP70-GFP detectable in the TIR field of illumination from synapse-associated ZAP70-GFP we conducted the following imaging sequence:

i. one 488 nm TIR image was acquired to quantitate freely diffusing and synapse-associated ZAP70-GFP. This was followed by:
ii. a 200 ms high-energy 488 nm TIR pulse to completely ablate the ZAP70-GFP signal in the TIR field of illumination and
iii. a second 488 nm TIR image separated from the bleach pulse by 0.8 seconds as a means to quantitate freely diffusing and rapidly recovering ZAP70-GFP, which was not associated with the plasma membrane yet detectable in TIR mode. This latter image (iii) was subtracted from the 488 nm image acquired prior to photobleaching (done in (i)) to quantitate synapse-associated ZAP70-GFP. Integrated and background-subtracted intensity values for ZAP70-GFP were divided by a constant (10^4^) and plotted against the number of receptor-engaged AF555-conjugated antigens per T-cell (see above).

### Fluorescence Recovery after Photobleaching (FRAP) analysis to determine the rate of ZAP70-GFP exchange at the immunological synapse

T-cells were seeded on the SLB for 5-10 minutes and then subjected to a TIR imaging protocol conducted in the following order:

i. 1 image of accumulated SLB-resident AF647-labeled antigen employing 640 nm laser excitation,
ii. 5 images of ZAP70-GFP each separated by 2 seconds employing 488 nm laser excitation and serving as internal pre-bleach control,
iii. a bleach pulse applying the 488 nm laser to ablate GFP fluorescence and conducted with the use of a slit aperture covering half the synaptic area, and
iv. 60 consecutive images of ZAP70-GFP acquired at 2 second time intervals using 488 nm laser excitation.

Integrated GFP fluorescence was measured for defined regions of interest (ROIs) after background subtraction and normalized by the average of the integrated intensity determined for the five pre-bleach control images. Recovery curves were fitted to a single exponential function to assess the time required for half of the maximal recovery.

### Quantitation of cell surface CAR or TCR molecules

CMVsp-ROR1-CAR-T-cells were stained with increasing concentrations of ROR1-AF647 or Alexa Fluor® 647 anti-human TCR α/β Antibody (Clone IP26, BioLegend). Mean fluorescence intensity (MFI) of each cell population was measured by FACS. The MFI of Quantitation beads (Quantum™ Alexa Fluor® 647 MESF, Bangs laboratories, INC) was also measured simultaneously to establish a calibration curve relating instrument channel values to standardized fluorescence intensity (MESF). Recorded MFI values of cell populations were then converted to MESF units following manufacturer’s instructions.

### Quantitation of synapse-associated phosphorylated ZAP70 and visualization of synaptic CD45

Approximately 10 minutes after seeding T-cells were fixed with PBS supplemented with 4% formaldehyde (Thermo Fisher Scientific), 1 mM Na_3_O_4_V and 50 mM NaF (both Sigma) for 20 minutes at room temperature. The fixation solution was then removed by rinsing the SLB-attached T-cells three times with washing buffer (PBS supplemented with 1 mM Na_3_O_4_V and 50 mM NaF). T-cells were permeabilized through incubation in PBS supplemented with 0.1% Triton™ X-100 Surfact-Amps™ (Thermo Fisher Scientific) for 1 minute. After rinsing three times with washing buffer supplemented with 3% BSA, the T-cells were incubated overnight at 4 °C with AF647-conjugated antibody reactive to ZAP70 phosphorylated at position 319 (Clone 1503310, BioLegend). The next day, unbound antibody was removed by rinsing the sample with washing buffer supplemented with 3% BSA. T-cells were then subjected to TIR microscopy as described above. Integrated intensity values of AF647-cojugated phosphorylated ZAP70 Ab image were divided by a constant (10^5^) and plotted as a function of SLB antigen density.

For visualizing synapse-associated CD45, the above-mentioned protocol was followed with the exception that cells were not permeabilized and were stained with Alexa Fluor 488 anti-human CD45 Antibody (Clone H130, BioLegend).

